# Retrotransposon proliferation coincident with the evolution of dioecy in *Asparagus*

**DOI:** 10.1101/048462

**Authors:** Alex Harkess, Francesco Mercati, Loredana Abbate, Michael McKain, J. Chris Pires, Tea Sala, Francesco Sunseri, Agostino Falavigna, Jim Leebens-Mack

## Abstract

Current phylogenetic sampling reveals that dioecy and an XY sex chromosome pair evolved once or possibly twice in the genus *Asparagus*. Although there appear to be some lineage-specific polyploidization events, the base chromosome number of 2n=2x=20 is relatively conserved across the *Asparagus* genus. Regardless, dioecious species tend to have larger genomes than hermaphroditic species. Here we test whether this genome size expansion in dioecious species is related to a polyploidization and subsequent chromosome fusion or retrotransposon proliferation in dioecious species. We first estimate genome sizes or use published values for four hermaphrodites and four dioecious species distributed across the phylogeny and show that dioecious species typically have larger genomes than hermaphroditic species. Utilizing a phylogenomic approach we find no evidence for ancient polyploidization contributing to increased genome sizes of sampled dioecious species. We do find support for an ancient whole genome duplication event predating the diversification of the *Asparagus* genus. Repetitive DNA content of the four hermaphroditic and four dioecious species was characterized based on randomly sampled whole genome shotgun sequencing and common elements were annotated. Across our broad phylogenetic sampling, *Ty-1 Copia* retroelements in particular have undergone a marked proliferation in dioecious species. In the absence of a detectable whole genome duplication event, retrotransposon proliferation is the most likely explanation for the precipitous increase in genome size in dioecious *Asparagus* species.

## Introduction

Fewer than 10% of flowering plant species are dioecious, the condition where individual plants are distinctly male or female (Ainsworth 2000). Gender in some dioecious plants can be governed by a sex chromosome pair, such as in papaya (*Carica papaya*), white campion (*Silene latifolia*), persimmon, *Rumex*, and garden asparagus (*Asparagus officinalis*) (Telgmann-Rauber *et al*. 2007; Ming *et al*. 2011; Akagi *et al*. 2014; Hough *et al*. 2014). The evolution of a distinct sex chromosome pair is hypothesized to be driven by the evolution of a non-recombining region between the X and Y (or Z and W) where tightly linked sex determination genes reside (Charlesworth and Charlesworth 1978). Given the repeated and independent evolution of dioecy across the angiosperm phylogeny, the transition from autosome to sex chromosome is undoubtedly governed by different sex determination genes and evolutionary processes, and consequently must be viewed in a taxon-specific context. Despite this diversity in sex chromosome evolution across the angiosperms, two particularly interesting associations can be seen in some dioecious systems coincident with variation in sexual system: the proliferation of repetitive elements and the occurrence of one or multiple whole genome duplication (polyploidy) events.

As a consequence of restricted recombination between regions of sex chromosomes, repetitive elements tend to persist and replicate in an unbalanced way, preferentially accumulating on hemizygous regions of Y and W chromosomes. Transposable elements can be broadly classified primarily by their means of transposition (Wicker *et al*. 2007); class I retrotransposons move by a “copy and paste” mechanism and replicate through an mRNA intermediate which ultimately results in a net increase of the element’s copy number, whereas class II transposable elements move through a DNA intermediate in a “cut and paste” fashion. Since Class I retrotransposons can range in length from 5 to 20 kilobases or longer, their proliferation can lead to drastic and rapid changes in genome size (Kidwell 2002). This accumulation, especially of active retroelements, can be clearly seen when comparing the relatively young papaya X and hermaphrodite-specific region of the Y (HSY) (VanBuren and Ming 2013). Unbalanced accumulation of transposons and other repetitive elements, paired with the inability for recombination to remove them along with other deleterious mutations, is likely a major factor that leads to the initial physical expansion and genic degeneration of a young, partially non-recombining Y or W chromosome (Steinemann and Steinemann 1998; Bachtrog *et al*. 2008; Bachtrog 2013). Transposons have also been directly implicated in the evolution of sex determination genes through disruption of gene expression. In melon (*Cucumis melo*), a class II hAT DNA transposon insertion is responsible for promoter hypermethylation and transcriptional repression of the zinc-finger transcription factor *CMWIP1*, heritably inducing the transition from monoecy to gynodioecy (Boualem *et al*. 2008).

An association between polyploidy and transitions in sexual system across the angiosperms is most clear in the *Fragaria* genus, where at least four independent whole genome duplication events have occurred across all major clades, leading to an abundance of polyploid dioecious species phylogenetically placed as sister to dioecious hermaphrodites (Rousseau-Gueutin *et al*. 2009; Ashman *et al*. 2013). However loss of dioecy has also been associated with an increase in ploidy, as seen in one clade of *Mercurialis* (Krähenbühl *et al*. 2002). The mechanisms that potentially relate whole genome duplication events to the evolution of sexually dimorphic populations are variable and poorly understood, though, again owing to the extreme complexity and species-specific nature of sex chromosome and dioecy.

Garden asparagus (*Asparagus officinalis L*.) is a particularly useful dioecious plant for studying sex chromosome evolution given that it has cytologically homomorphic X and Y sex chromosomes, suggesting that the transition from hermaphroditism to dioecy was recent (Telgmann-Rauber *et al*. 2007; Kubota *et al*. 2012). Coincident with the evolution of dioecy was a range shift from South Africa into North Africa, Europe, and Asia (Štajner *et al*. 2002; Kubota *et al*. 2012; Norup *et al*. 2015). It was previously reported that dioecious *Asparagus* species tend to have larger genomes than hermaphrodites, but there was no evidence supporting a whole genome duplication event that separates the dioecious species from the hermaphrodites (Kuhl *et al*. 2005). The base chromosome number of 2n=2x=20 is generally consistent across the genus except for instances of very recent polyploidization in some species (Kanno and Yokoyama 2011). These findings suggest one of two hypotheses may be responsible for an increase in genome size: one possibility is that a whole genome duplication occurred in the last common ancestor of all dioecious species, followed by a chromosome fusion or reduction, and another possibility is that repetitive DNA has proliferated to drive the increase in the genome sizes of dioecious species. Here, we test both hypotheses by first leveraging transcriptome assemblies for one hermaphroditic and one dioecious species to identify the relative timing of whole genome duplication events in the genus *Asparagus*. We then use shallow Roche 454 whole genome shotgun sequencing from four hermaphrodites and four dioecious species that are sampled from across the phylogeny to assess the repetitive element content of each species in relation to its genome size.

## Results and Discussion

### Genome size increases in dioecious Asparagus

Genome sizes and ploidy vary greatly across the order Asparagales, with 1C values ranging from 0.3-88.2 pg (Leitch *et al*. 2010). Diploid dioecious *Asparagus* species have been reported as having genome sizes nearly double the size of diploid hermaphroditic congeners (Štajner *et al*. 2002, Fukuda et al. 2005, Kubota et al., 2012). We first confirmed this by generating or supplementing published genome size estimations for eight *Asparagus* species, four hermaphrodites and four dioecious species, sampled across all major clades of the *Asparagus* phylogeny (Kubota *et al*. 2012) (Table 1). All individuals have been documented as diploids (2n=2x=20) except for *A. maritimus*, a hexaploid (Štajner *et al*. 2002; Kanno and Yokoyama 2011). Flow cytometry-derived genome sizes (pg/1C) for hermaphrodites range from 0.72 to 1.06, while dioecious species range from 1.09 to 1.37. Dioecious species tend to have larger genome sizes than hermaphroditic species (Unpaired *t* test, p = 0.0173). An outlier is the hermaphrodite *Asparagus asparagoides* with a relatively large genome size (1C = 2.40; Dixon’s Q test, p = 0.074).

**Table 1:** Genome sizes, 454 pyrosequencing and repetitive element clustering

| Species | Sexual System | pg/nucleus<br>(mean ± SD) | 1C<br>value | Raw<br>Reads | Filtered<br>Reads | Clustered reads<br>(%) |
| --- | --- | --- | --- | --- | --- | --- |
| **A. officinalis** | Dioecious | 2.74 ± 0.044 | 1.37 | 29,677 | 26,525 | 54.4% |
| **A. maritimus** | Dioecious | 7.87 ± 0.204<sup>a</sup> | 1.31 | 49,616 | 45,036 | 53.7% |
| **A. aphyllus** | Dioecious | 2.49 ± 0.007 | 1.25 | 47,322 | 42,808 | 58.9% |
| **A. stipularis** | Dioecious | 2.17 ± 0.005 | 1.09 | 30,405 | 27,911 | 56.4% |
| **A. falcatus** | Hermaphrodite | 2.11 ± 0.007 | 1.06 | 26,836 | 24,304 | 60.4% |
| **A. virgatus** | Hermaphrodite | 1.66 ± 0.055 | 0.83 | 45,043 | 41,053 | 45.0% |
| **A. pyramidalis** | Hermaphrodite | 1.44 ± 0.037<sup>a</sup> | 0.72 | 56,197 | 51,293 | 53.8% |
| **A. asparagoides** | Hermaphrodite | 4.80 ± 0.062 | 2.40 | 41,952 | 37,435 | 59.2% |
| **Sum** |  |  |  | 247,755 | 224,804 |  |
| **Average** |  |  |  | 41,293 | 37,467 |  |

<sup>a</sup>Data from Stajner et al. (2002)

#### No evidence for a dioecy-specific polyploidy event

We employed a phylogenomics approach to test whether a whole genome duplication event separates the dioecious and hermaphroditic species in *Asparagus*. Transcriptome assemblies were generated for two species sampled broadly across the phylogeny: a basal diploid hermaphrodite (*A. asparagoides*; 2n=2x=20) and diploid dioecious garden asparagus (*A. officinalis*; 2n=2x=20). Intraspecific paralog pairs and interspecific orthologous gene pairs were inferred to generate *Ks* (synonymous substitution rate) distributions and assess the relative timing of whole genome duplication event relative to speciation events (Blanc and Wolfe 2004, Cui et al. 2006, McKain et al. 2012, Doyle and Egan 2010). Despite being an outlier in terms of genome size, *A. asparagoides* was utilized for the comparison given that it is a basal member of the genus, shares the same diploid chromosome count as *A. officinalis*, and that transcriptome-based *Ks* analyses are independent of genome size.

Transcriptome assembly and translation results for the two species are presented in Supplementary Table 1. One distinct, shared polyploidization event (*Ks* ~ 0.5) was inferred from the *Ks* frequency distribution of paralogous pairs in both *Asparagus* species (Figure 1; Supplementary Table 2). Additionally, orthologous pairs exhibit a *Ks* peak close to 0, representing low divergence and suggestive of recent diversification of species and/or similar mutation rates. Comparison of orthologs and paralogs demonstrates that at least one detectable genome duplication event occurred before the diversification of the *Asparagus* genus (Figure 1).

**Figure 1:**
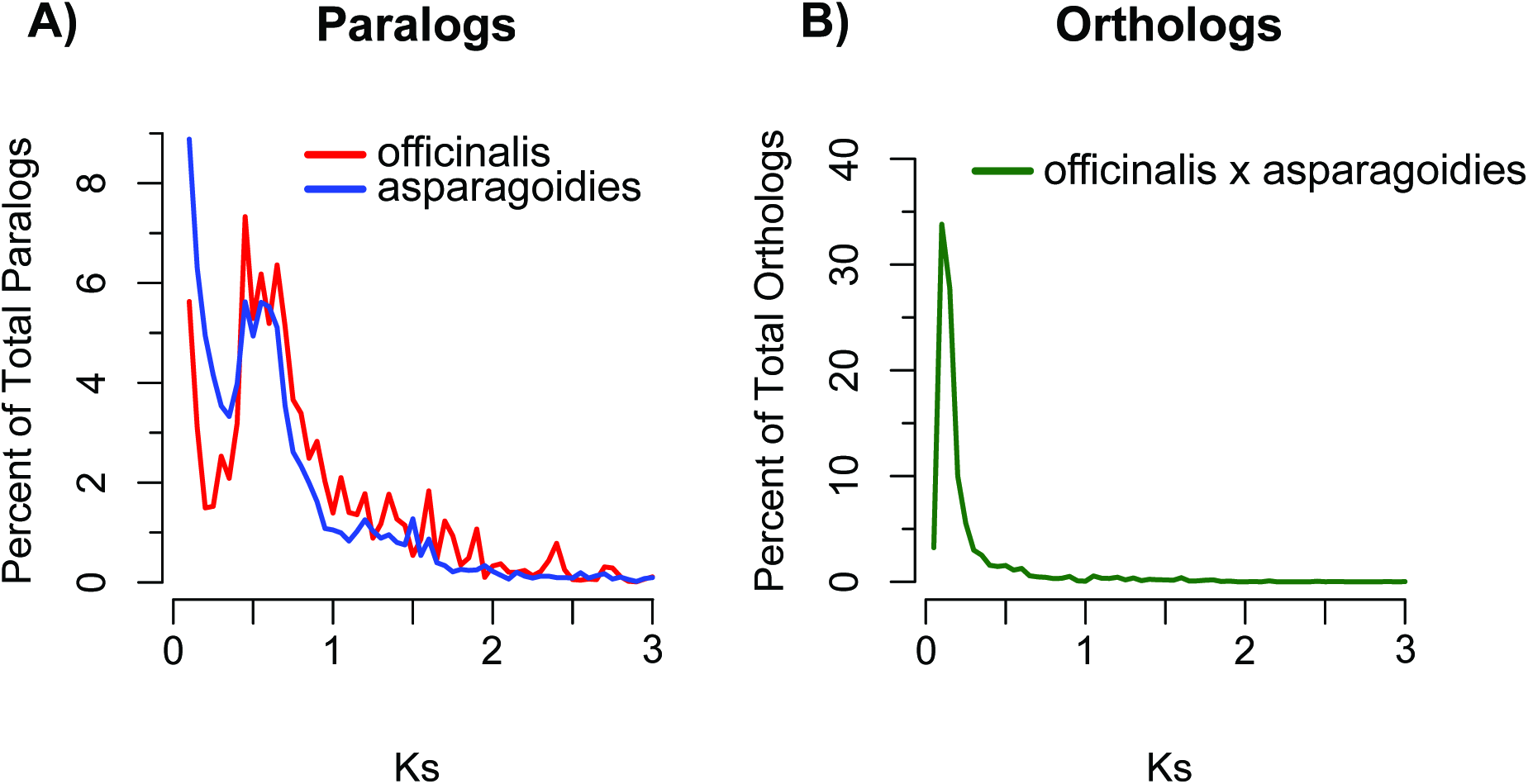
Transcriptome-based *Ks* frequency distributions for A) paralogous and B) orthologous pairs of dioecious *A. officinalis* and hermaphroditic *A. asparagoides*. Paralogous and orthologous *Ks* distributions suggest a shared whole genome duplication event at *Ks* ~ 0.5 that occurred before the diversification of the *Asparagus* genus.

A major limitation with *Ks* analyses is that more recent duplication events are difficult to detect (Blanc and Wolfe 2004; Cui et al. 2006). This issue is exacerbated when using *de novo* transcriptome assemblies, where recently duplicated paralogs can be computationally mistaken as alleles and incorrectly collapsed into a single transcript sequence during the assembly process. Given that there are no current age estimates for the divergence of the *Asparagus* lineage, we cannot exclude the possibility that a more recent duplication event, such as one that may have co-occurred with the evolution of dioecy, could be undetectable by transcriptome data. However, such a whole genome duplication event would need to be a followed by multiple chromosome fusion or loss events to reduce the chromosome number back to 2n=2x=20 found in most karyotyped dioecious *Asparagus* species (Kanno and Yokoyama 2011). Taken together, the overlapping paralogous and orthologous *Ks* distributions do not support the hypothesis that a whole genome duplication event occurred coincident with the evolution of dioecy.

#### Lineage-specific expansion of transposable elements

Given the lack of evidence that ancient polyploidy was responsible for the larger genome sizes of dioecious *Asparagus* species relative to hermaphroditic species, we assessed the alternative hypothesis that the genome size increases in dioecious species was due to transposon amplification. We utilized whole genome shotgun sequence reads to assess the repetitive content of hermaphrodite and dioecious *Asparagus* species using the RepeatExplorer Galaxy server (http://www.repeatexplorer.org). Briefly, this method utilizes all-by-all read comparisons followed by Louvain clustering (Blondel *et al*. 2008) to place reads into unbiased clusters of putative high copy elements, followed by a RepeatMasker annotation and cap3 assembly (Huang and Madan 1999).

A total of 327,048 raw reads were sequenced for the eight genomes using Roche 454 FLX chemistry, with genome coverages ranging from 0.0051X to 0.0234X (Supplementary Table 3). After removing duplicate reads that were likely clonal, 321,865 total reads remained for analysis. To improve clustering, we then removed reads less than 100nt long, yielding a filtered set of 296,365 reads (mean = 37,047 reads per species) with a mean length of 321nt. The filtered set of reads was concatenated and clustered using the RepeatExplorer pipeline, placing 162,435 reads into 29,643 repetitive element clusters (Table 1). Repetitive element clusters were filtered by read count, requiring at least 0.01% of the total filtered reads (30 reads), amounting to 336 clusters for downstream analysis. These clusters were annotated against a custom RepeatMasker database generated with additional data for dioecious *A. officinalis*. For a given cluster of repetitive elements, the repetitive fraction of each species’ genome was calculated as the number of a given species’ reads in a cluster divided by the total number of reads sequenced for that species, represented as a percentage.

Multidimensional Scaling (MDS) analysis of the genomic proportions for all clusters shows that dioecious and hermaphroditic species form two distinct clusters (Figure 2). In general, *Gypsy* and *Copia* retrotransposons dominate the genomic landscape for all sampled *Asparagus* species (Figure 3). In all four dioecious species, *Gypsy* retrotransposons occupy a larger percentage of each genome than in the hermaphrodites, although *Copia* elements have distinctly expanded in the dioecious species (Figure 2). This suggests that both *Gypsy* and *Copia* elements increased in copy number in the dioecious species, and the proliferation of *Copia* elements was a more substantial contributor to the expansion of dioecious genome sizes.

**Figure 2:**
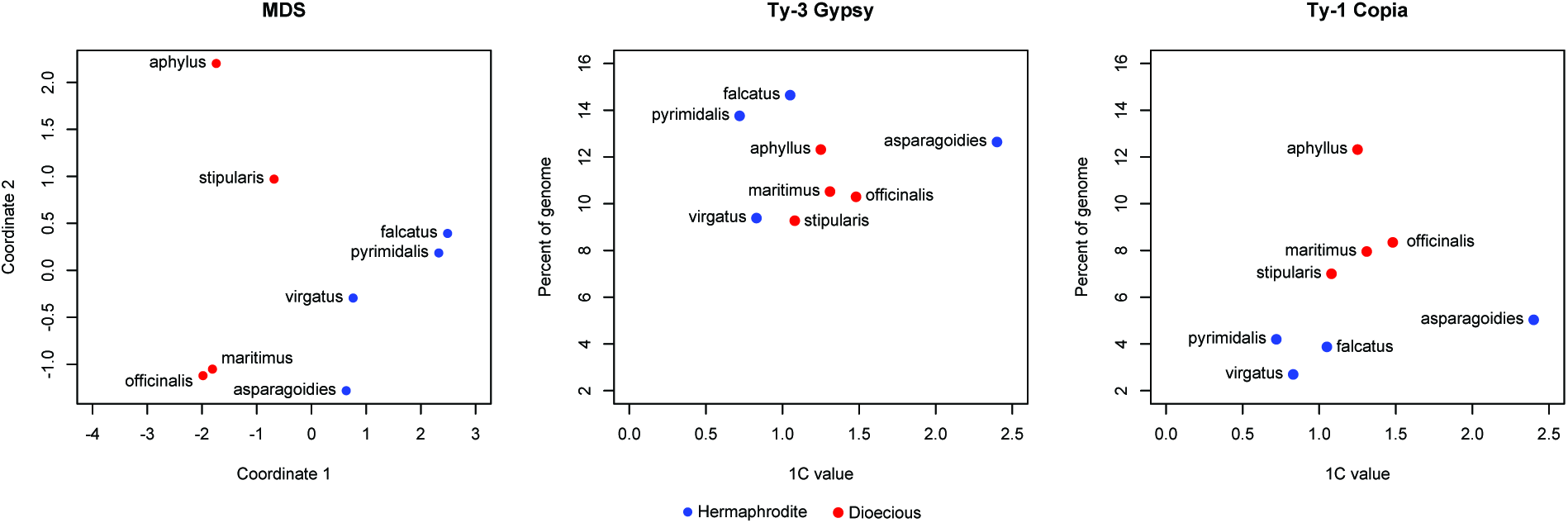
Multidimensional scaling (MDS) and relationship of genome size to *Gypsy* and *Copia* retroelement content for both dioecious and hermaphroditic genomes. Blue dots represent hermaphroditic species while red dots represent dioecious species.

**Figure 3:**
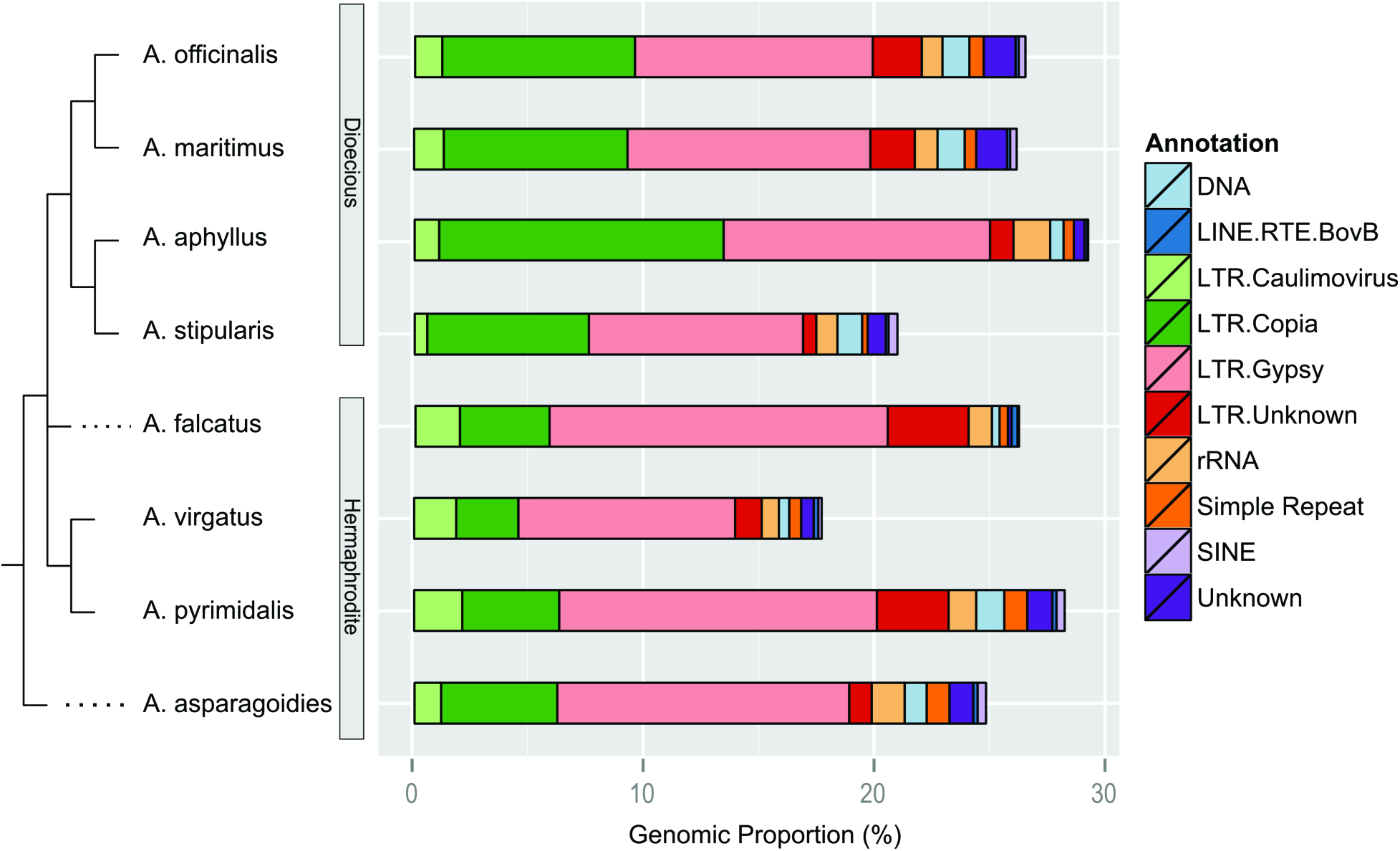
Cladogram of *Asparagus* species relationships with high copy repetitive elements. High copy elements refer to clusters with greater than 0.01% of the total read count in the multispecies analysis, able to be most confidently annotated against the custom *A. officinalis* repetitive element database. DNA transposons from several families were collapsed into a single annotation class.

We identified 46 repetitive element clusters that were private to the dioecious species and 37 clusters that were private to all hermaphroditic species. In the dioecious species, 26 clusters were *Gypsy* and 7 clusters were *Copia*, whereas in the hermaphroditic species, 12 clusters were *Gypsy* and 11 clusters were *Copia*. This suggests that there is active turnover of transposable elements in the *Asparagus* genus, perhaps coincident with the evolution of dioecy and a sex chromosome. Additionally, it is possible that a small number of *Copia* elements may be largely responsible for the genome size expansion in dioecious species, but this would require whole genome assemblies and annotations as RepeatExplorer is limited in ability to finely delimit elements.

One caveat for performing a single repeat clustering analysis including all species (as opposed to individually analyzing each species) is that low frequency or moderately diverged sequences from phylogenetically distant species may not cluster. Additionally, there could be less power for detecting species-specific transposon family proliferations. Consequently, these estimates of repetitive element content are certainly underestimates of the total proportion of repetitive element content in each species’ genome. To understand the level of difference in these two analysis types, we generated 893,623 additional 454 shotgun reads (mean length 526nt) for a mature double haploid YY *A. officinalis* individual and ran the RepeatExplorer pipeline with this single species. The repeat content was estimated at 71.1%, much greater than the 54.4% that was estimated by concatenating eight species in a single analysis. This result suggests that the genomic proportions of transposons estimated through multispecies read clustering in this study should be interpreted as being underestimates, biased towards high copy elements with lower divergence between species, and used mostly for comparisons of high copy element percentages between species. The advantage of this analysis is that direct comparisons for a given transposon cluster can be assessed across all species, without the need to perform additional clustering between species.

The method of repeat quantification and sequence read type also largely affects the estimated proportion of repetitive elements. Repetitive element content has previously been estimated for *A. officinalis* in at least three separate studies. Vitte *et al*. (2013) directly annotated garden asparagus Bacterial Artificial Chromosome (BAC) assemblies for transposon content. By comparing the sequence alignment identity of intact LTRs from retroelements and applying a clock estimation from rice retroelement divergence (Ma and Bennetzen 2004), Vitte *et al*. estimated that the majority of the asparagus genome is comprised of young, recently inserted (< 6 million years ago) and nested retroelements. Li *et al*. (2014) took a high-throughput sequencing approach and inferred that the garden asparagus genome is 53% repetitive by *de novo* assembling genomic paired end 100nt Illumina reads into a ~400Mbp assembly with a scaffold N50 of 1504nt. Hertweck (2013) took a similar approach with 80 bp Illumina read data and independently estimated 47% of the garden asparagus genome as comprising repetitive elements. We hypothesize that our much higher estimation of 71.1% repetitive content is largely due to the increased detection power coming with longer 454 reads relative to 80-100 bp Illumina reads and our use of RepeatExplorer’s unique assembly-free, graph-based clustering and annotation of individual long reads.

#### Transposon clustering yields phylogenetic signal

Clustering of the genomic proportions for the 100 largest *Gypsy* and *Copia* retrotransposon clusters also reveals phylogenetic signal in the data (Figure 4). The deepest branch divides the hermaphroditic and dioecious species from each other, and all species are paired with their closest phylogenetic neighbor given the current phylogeny and sampling from Kubota *et al*. (2012), with exception for the earliest diverging species in the genus, *A. asparagoides*. The genomic proportions of repetitive elements have been used to identify phylogenetic signal in several plant species with species relationships that have been difficult to resolve with traditional low copy gene sequencing (Dodsworth *et al*. 2014). While our clustering approach may be less able to detect low and medium-frequency repeats compared to Dodsworth *et al*. (2014), here we show a complementary analysis that yields similar results using high copy transposon clusters.

**Figure 4:**
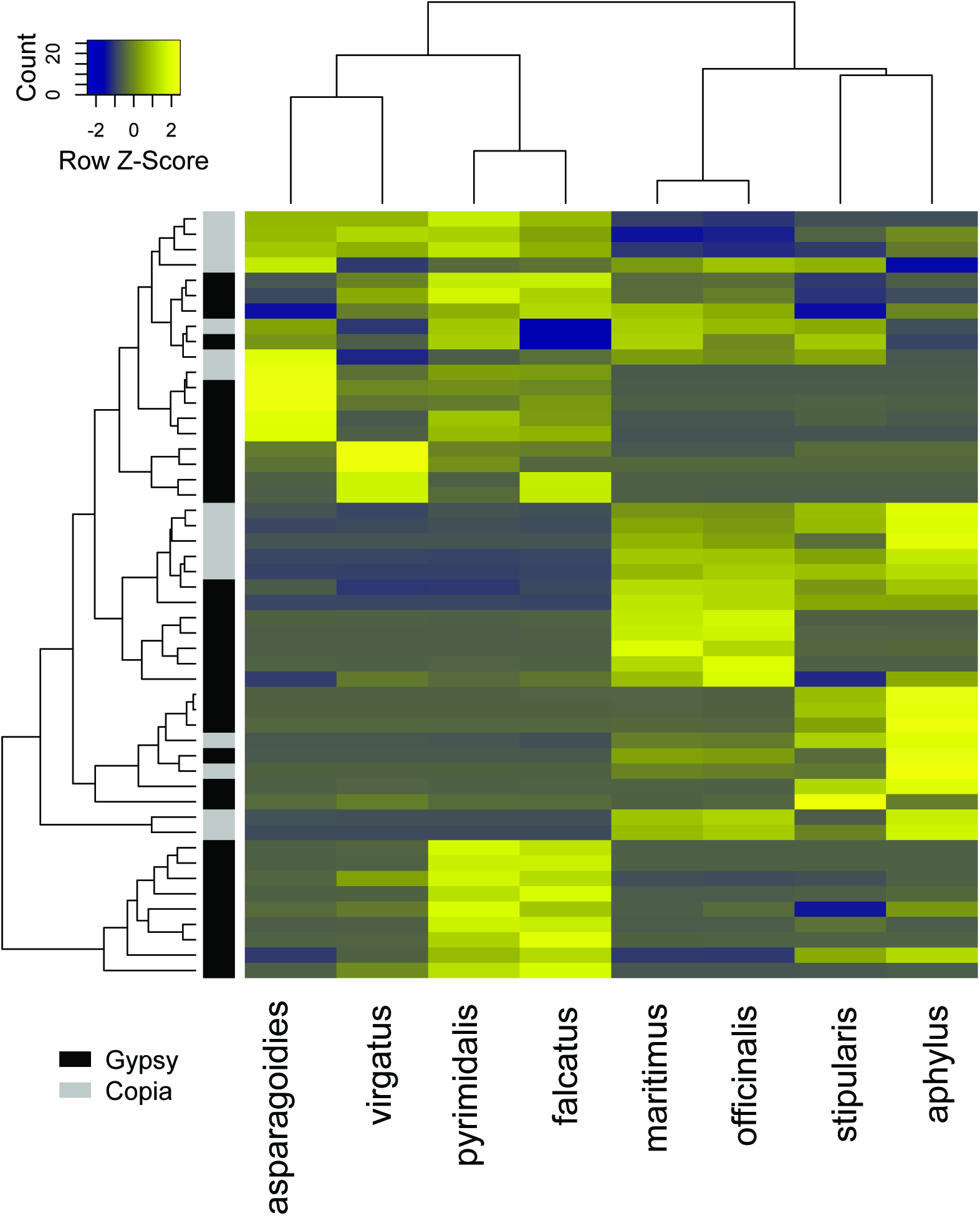
Heatmap clustering of 100 largest *Gypsy* or *Copia* element clusters. Rows represent individual clusters, annotated as *Gypsy* (black) and *Copia* (grey).

Recently, Norup et al. (2015) proposed two origins of dioecy within *Asparagus*, an alternative to the previously hypothesized single origin (Kubota *et al*. 2012). Our sampling includes species derived from both of the hypothesized origins of dioecy from Norup et al. (2015), which indicates that dioecy evolved in one clade that includes *A. officinalis* and *A. maritimus*, as well as another clade that includes *A. stipularis* and *A. aphyllus*. In the case of multiple origins of dioecy, without hermaphroditic outgroup species for each origin, our limited sampling does not allow us to describe the potentially different repetitive element radiations in the two dioecious clades. Further, it is possible that transposon proliferation and genome size increase occured in the common ancestor of both dioecious lineages, predating the origin of dioecy. Rigorous testing of a general relationship between transposon activity and the origin of sex chromosomes will come with future meta-analyses including data from this and other comparative studies of transposon activity in hermaphrodite and dioecious lineages

Several mechanisms exist to remove repetitive element DNA from obese, transposon-dense genomes. One mechanism is the formation of small chromosomal deletions by illegitimate recombination, which usually occurs by slip-strand mispairing or non-homologous end joining (NHEJ) (Hawkins *et al*. 2009). Another mechanism is unequal intra-strand homologous recombination between the directly repeated LTRs of retrotransposons, resulting in a solo LTR remaining (Devos *et al*. 2002). The half-life for retrotransposon occupancy seems to be relatively short in several plant species (Ma *et al*. 2004; Wang and Liu 2008; Charles *et al*. 2008), suggesting that these removal mechanisms are actively purging transposons. In most angiosperm genomes, *Gypsy* and *Copia*-type retrolements typically dominate the repetitive landscape. Extreme cases of transposon purging and genome compression are evidenced by the *Utricularia gibba* genome, comprising only about 3% transposable elements (Ibarra-Laclette *et al*. 2013). Compared to *Asparagus*, similar cases of lineage-specific transposon expansion have been found in the Asteraceae, where a small number of *Gypsy* families have been expanding since the branch leading to the Asteraceae (Staton and Burke 2015). We hypothesize that the proliferation of both *Gypsy* and *Copia* retroelements in dioecious lineages is associated with two coincident events in Asparagus evolution: range expansion and the origin of dioecy. As others have documented, range expansion out of South Africa is associated with a transition of ancestrally hermaphroditic *Asparagus* species to dioecy within a clade distributed across Europe and Asia. (Štajner *et al*. 2002; Kuhl *et al*. 2005; Kanno and Yokoyama 2011; Kubota *et al*. 2012; Norup *et al*. 2015). Founder populations formed during this range expansion with small effective population sizes may have been especially susceptible to weakly deleterious transposon proliferation due to the reduced strength of purifying selection relative to populations with large effective sizes (Lynch *et al*. 2011). In addition, the origin of sex chromosomes alone may have promoted proliferation of retrotransposons. Suppressed recombination within the region of the sex chromosomes where gender determination genes reside in the first dioecious *Asparagus* species may have harbored active retrotransposons. Young and old plant Y chromosomes in *Silene* and papaya can be replete with or entirely missing tandem arrays and LTR retroelements that distinguish them from both the X and other autosomes (Pritham *et al*. 2003; Filatov *et al*. 2009; VanBuren and Ming 2013). Recombination is selected against in these regions of a sex chromosome given that recombination could break apart genes influencing male and female function, leading to the formation of neuters. This selection on young sex chromosomes may drive the maintenance and proliferation of LTR retrotransposons, which in concert with faster mutation rates and background selection may lead to the initial expansion and subsequent degeneration of sex chromosomes (Engelstädter 2008).

## Methods

### Flow cytometry genome size estimation

The genome sizes of *A. officinalis*, *A. virgatus* and *A. asparagoides* were estimated by flow cytometry at the Benaroya Research Institute at Virginia Mason in Seattle, Washington. Nuclei isolations from a single mature leaf were analyzed in three technical replicates for each species. The genome sizes of *A. aphyllus*, *A. stipularis*, and *A. falcatus* were estimated by flow cytometry using the known genome size of *A. officinalis* (1C-value = 1.37 pg) as a reference standard. Ten plants for each species, grown in greenhouse, were sampled and three randomly selected plants were analysed. The analysis was carried out with the Partec PAS flow cytometer (Partec, http://www.partec.de/), equipped with a mercury lamp. Fully expanded leaves (0.1 g) were chopped in 300 μl nuclei extraction buffer (CyStain ultraviolet Precise P Nuclei Extraction Buffer; Partec, Münster, Germany) for 30-40 s. The solution was filtered through a 30 mm CellTrics disposable filter (Partec), and 1.2 ml of staining solution containing 4,6-diamidino-2-phenylindole was added. The relative fluorescence intensity of stained nuclei was measured on a linear scale and 4,000-5,000 nuclei for each sample were analysed (Galbraith et al., 1998). DNA content histograms were generated using the Partec software package (FloMax). Given that the X and Y chromosomes in garden asparagus (*Asparagus officinalis*) are cytologically homomorphic (Deng *et al*. 2012) representing a lack of degeneration and the relatively young age of the Y, we did not discern between potential sex differences in the dioecious species.

### Transcriptome-based Ks analysis

Transcriptomes from dioecious *A. officinalis* and hermaphroditic *A. asparagoides* were used to infer a putative whole genome duplication event in the genus *Asparagus*. The transcriptome assembly and translation for *A. officinalis* was taken from Harkess *et al*. (2015) (http://datadryad.org/resource/doi:10.5061/dryad.92c60). We generated leaf RNA-Seq for *A. asparagoides* by first isolating total RNA from mature leaf tissue using a Qiagen RNeasy Plant Mini kit. Total RNA quantity and quality was assessed using an RNA Nano chip on the Bioanalyzer 2100. A sequencing library was generated using the TruSeq RNA Library Prep Kit v2 (Illumina) according to manufacturer’s instructions using 1ug of total RNA input. The library was sequenced with paired end 100nt reads on an Illumina HiSeq2000, generating 55,686,513 read pairs (nearly 11 gigabases of data). Reads were quality trimmed using Trimmomatic (v0.32), removing sequencing adapters and clipping 3’ and 5’ read ends with a quality score lower than Phred 5. Cleaned reads were assembled using Trinity (r20140717) with default parameters. We filtered transcript isoforms with low support by removing isoforms with less than 0.01% of the Trinity gene subcomponent read support. Coding sequence and peptide translations were inferred using TransDecoder (r20140704) with default settings. Raw sequence reads for *A. asparagoides* has been deposited under BioProject (ID here after acceptance).

Using a pipeline from McKain et al. (https://github.com/mrmckain/FASTKs), we first identified putative paralogs in each filtered transcriptome assembly using all vs. all blastn (1e-40 cutoff). Peptide sequences for hit pairs longer than 100 amino acids were aligned using MUSCLE (v3.8.31), then codon alignments were inferred using PAL2NAL (v13) (Suyama *et al*. 2006). For each paralog pair, *Ks* was calculated using CodeML in PAML (Yang 2007) (v4.8).

### 454pyrosequencing and transposon quantification

Whole genomic DNA was extracted from four hermaphroditic and four dioecious species using a CTAB method (Doyle and Doyle 1987). Sequencing libraries were prepared using the Roche 454 GS FLX Titanium library preparation kit according to manufacturer instructions. Raw reads were first de-duplicated to remove probable emulsion PCR sequencing artifacts, then filtered to remove reads less than 100nt long. Read names from all species were first prepended with a unique species identifier and concatenated. The RepeatExplorer (v0.9.7.4) pipeline (http://www.repeatexplorer.org) was then used to cluster, assemble, and annotate all filtered shotgun reads against a custom garden asparagus RepeatMasker database (see below) using otherwise default settings. Clustering and heatmap production of the 100 largest transposon clusters was performed using heatmap.2 in the gplot package in R (v3.2.1) using default settings; a distance matrix was generated using Euclidean distances, and hierarchical clustering was performed using “complete” clustering.

To improve the annotations of repetitive element clusters generated through the RepeatExplorer pipeline instead of utilizing default RepeatMasker libraries, we generated a much higher coverage of 454 reads for *A. officinalis* to build a comprehensive database of annotated exemplar repeats for the *Asparagus* genus. A custom garden asparagus RepeatMasker database was generated using similar methodology. A total of 893,623 454 FLX Titanium reads were generated from leaf tissue of a doubled haploid (YY) garden asparagus individual. Reads were more stringently filtered to a 150nt minimum length. The same version of RepeatExplorer was then run, and the resulting cap3 consensus assemblies for each cluster were annotated using RepeatClassifier, part of the RepeatModeler (v1.0.8) suite, with default settings. A total of 22,361 sequences greater than 150nt in length and with annotations were retained for annotating all repetitive element clusters and are available at (**DRYAD LINK)**. Raw 454 shotgun sequence data for all individuals have also been deposited in Dryad.

## Data availability

Raw RNA-Seq reads for *A. officinalis* will be deposited in SRA upon acceptance. Raw 454 shotgun reads will be deposited in Dryad upon acceptance. Additionally, the Dryad repository will contain the custom *A. officinalis* repetitive element database.

